# Mapping RNA-Binding Proteins on the Ribosome by Tethered Micrococcal Nuclease

**DOI:** 10.1101/2025.10.24.684434

**Authors:** Chia Yi Yao, Simpson Joseph

## Abstract

RNA-binding proteins (RBPs) are essential regulators of post-transcriptional gene expression, influencing mRNA processing, translation, and stability. Defining their binding sites on RNA is key to understanding how they assemble into functional ribonucleoprotein (RNP) complexes, but existing footprinting and crosslinking approaches often yield low signal-to-noise, variable efficiency, or require highly purified complexes. To address these limitations, we developed Tethered Micrococcal Nuclease Mapping (TM-map), a sequencing-based strategy that determines the three-dimensional binding sites of RBPs on RNA. In TM-map, the RBP is fused to micrococcal nuclease (MNase), which upon Ca²⁺ activation cleaves proximal RNA regions, producing fragments whose 3′ termini report the spatial proximity of the fusion. We first validated TM-map using the bacteriophage MS2 coat protein bound to its cognate RNA stem-loop engineered into the *Escherichia coli* ribosome. Cleavage sites mapped within ∼15 Å of the stem-loop, confirming that tethered MNase accurately reports local structure on the ribosome surface. We then applied TM-map to the *Drosophila* Fragile X Mental Retardation Protein (FMRP), a translational regulator with an unresolved ribosome-binding site. Both N-and C-terminal MNase-FMRP fusions produced reproducible cleavage clusters on the 18S rRNA localized to the body and head of the 40S subunit. The similar profiles suggest that FMRP’s termini are conformationally flexible and sample multiple orientations relative to the ribosome, consistent with a dynamic interaction rather than a fixed binding mode. TM-map thus provides a simple, high-resolution, and generalizable approach for visualizing RBP-RNA interactions within native RNP assemblies.

**Table of Contents Graphic:** 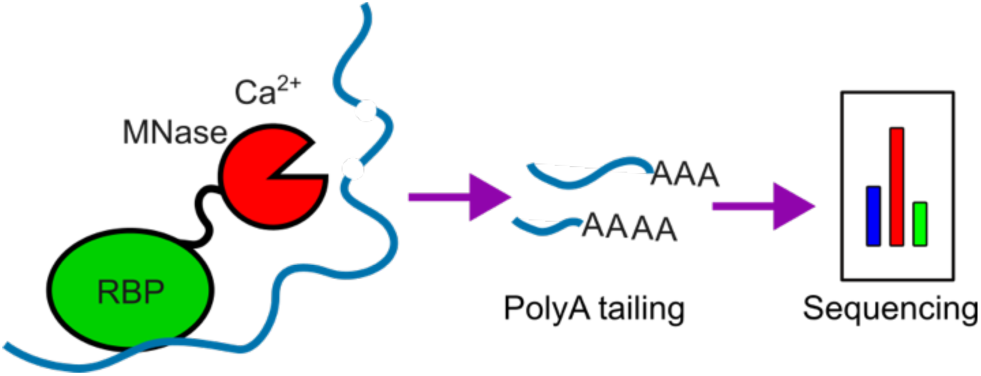

## INTRODUCTION

RNA-binding proteins (RBPs) play central roles in post-transcriptional gene regulation and RNA metabolism. They function as structural components of ribonucleoprotein (RNP) assemblies such as the ribosome, spliceosome, and telomerase, and participate in essential processes including mRNA biogenesis, transport, and localization. ^1–4^ RBPs also directly modulate gene expression by associating with the ribosome, forming part of the miRNA-mediated RNA interference complex, or interacting with long non-coding RNAs. ^1–4^ Given their broad regulatory roles, mutations that disrupt RBP function are implicated in neurological disorders, cancer, and other diseases. ^5–9^ Thus, defining RNA-protein interactions is fundamental for understanding the molecular basis of development, physiology, and disease.

A variety of biochemical and structural approaches have been developed to identify RBP-binding sites on RNA. Classical enzymatic and chemical footprinting methods, crosslinking-based approaches, and directed hydroxyl radical probing have each contributed to our understanding of RNA-protein interfaces. ^10–16^ In recent years, many of these methods have been adapted for transcriptome-wide analysis using high-throughput sequencing, enabling in vivo identification of RBP-binding sites. ^12,17–19^ However, these approaches each have notable limitations. Enzymatic and chemical footprinting often yield low signal-to-noise ratios, complicating the precise localization of the binding site. ^4,10,20,21^ Crosslinking-based methods suffer from low and variable crosslinking efficiency, which can lead to incomplete or ambiguous mapping of RNA-protein interactions. ^22^

To overcome these challenges, we sought to develop a method that directly couples the physical proximity of an RNA-binding protein (RBP) to its target RNA with a quantitative, sequence-based readout. Our goal was to create an approach that is experimentally straightforward, yields high signal-to-noise, and can be broadly applied to diverse RNA-protein assemblies. This concept was inspired by pioneering work employing Micrococcal Nuclease (MNase) fusions to map protein-nucleic acid interactions. Henikoff and colleagues introduced the “Cut & Run” method to identify transcription factor binding sites on DNA by activating Ca²⁺-dependent MNase to cleave adjacent DNA and recover the resulting fragments for sequencing. Similarly, Milkereit and co-workers fused MNase to specific ribosomal proteins in *Saccharomyces cerevisiae* to probe the local three-dimensional architecture of rRNA within ribosomal subunits.^23^ More recently, MNase fusions to ribosomal proteins and ribosome assembly factors have been used to interrogate the spatial organization of rRNA during ribosome maturation. ^24,25^ Together, these studies established MNase fusions as versatile molecular reporters that translate spatial proximity into precise, sequence-defined cleavage patterns.

Building on this concept, we developed a technique - Tethered MNase Mapping (TM-map) - to identify the binding sites of RBPs on RNA and within large RNP complexes. TM-map employs a fusion of MNase to the protein of interest, which, upon activation by Ca²⁺, cleaves nearby RNA, generating a precise signature of its spatial proximity. The resulting fragments are poly(A)-tailed, sequenced, and computationally mapped to determine the position of MNase cleavage at nucleotide resolution. As proof of principle, we first tested the method using the bacteriophage MS2 coat protein bound to its cognate RNA stem-loop engineered into the *E. coli* ribosome. ^26–28^ The high-affinity and well-characterized MS2-stem-loop interaction, together with an available cryo-EM structure, provided an ideal benchmark to validate TM-map. ^29,30^

We then extended TM-map to study the binding of *Drosophila* Fragile X Mental Retardation Protein (FMRP) on the 80S ribosome. Previous biochemical and cryo-EM studies have shown that FMRP associates with the ribosome and inhibits translation, but its precise binding position remains unclear due to the limited resolution of the existing structure. ^31–36^ Using MNase fusions at both the N-and C-termini of FMRP, we mapped cleavage sites on *Drosophila* rRNAs to infer the three-dimensional position of FMRP on the ribosome. Beyond validating known structural features, this approach provides a scalable framework for studying RBP-RNA interactions within diverse macromolecular assemblies where direct structural visualization remains challenging.

## MATERIALS AND METHODS

### GST-MS2-MNase Construct Design and Protein Purification

The *GST-MS2-MNase* fusion construct was generated by inserting the *MNase* gene (Addgene # 123461), preceded by a six-amino acid flexible linker, into a plasmid encoding *GST-MS2* using homologous recombination. ^28^ The resulting plasmid (*pGST-MS2-MNase*) was verified by complete plasmid sequencing.

For protein expression, *E. coli* BL21(DE3) cells were transformed with *pGST-MS2-MNase*. Transformants were cultured overnight at 37 °C in LB medium containing 100 µg mL⁻¹ ampicillin. Each 1 L of LB medium supplemented with 100 µg mL⁻¹ ampicillin was inoculated with 5 mL of the overnight culture and grown at 37 °C until the optical density at 600 nm reached 0.6. Protein expression was induced with 0.1 mM IPTG, and cultures were incubated for 18 h at room temperature. Cells were harvested by centrifugation and resuspended in lysis buffer (20 mM Tris-HCl, pH 7.4; 150 mM KCl; 2 mM DTT; 5% v/v glycerol). Cell disruption was performed using a microfluidizer (18,000 psi, two passes), and the lysate was clarified by two sequential centrifugations in a Ti-70 rotor (13,000 rpm, 30 min, 4 °C).

The supernatant was incubated with glutathione-Sepharose FF resin (10 column volumes, pre-equilibrated) for 30 min at 4 °C with gentle mixing. The resin was packed into a gravity column, washed with 100 column volumes of PBS containing 2 mM DTT, and the bound protein was eluted with 50 mM Tris-HCl (pH 8.0) containing 10 mM reduced glutathione. Eluted fractions were dialyzed against 3 × 1 L PBS supplemented with 50% (v/v) glycerol using a 6 - 8 kDa molecular-weight-cutoff membrane. Protein concentration was determined using the Bradford assay, and aliquots were stored at-20 °C.

### Plasmid Replacement and Purification of 70S-MS2-SL Ribosomes

The *70S-MS2-SL* construct was transformed into *E. coli* strain SQZ10 (Δ7 *rrn*) harboring the helper plasmid *pHK-rrnC + sacB* (kanamycin resistant) ^37^. Transformants were cultured overnight at 37 °C in LB medium containing 100 µg mL⁻¹ ampicillin. To select for plasmid replacement, overnight cultures were plated on 2YT agar supplemented with 8% (w/v) sucrose and 100 µg mL⁻¹ ampicillin. Colonies were screened for kanamycin sensitivity by replica plating, and confirmed clones were used for large-scale ribosome purification.

Starter cultures (4 × 5 mL LB + 50 µg mL⁻¹ kanamycin) were grown overnight at 37 °C and used to inoculate six 500 mL LB cultures containing 50 µg mL⁻¹ kanamycin. Cultures were grown at 37 °C to an OD₆₀₀ of 0.5, harvested by centrifugation, and resuspended in lysis buffer composed of 50 mM Tris-HCl (pH 7.4), 10 mM MgCl₂, 100 mM NH₄Cl, 6 mM β-mercaptoethanol, and 0.5 mM EDTA. The suspension was pelleted again (8,000 rpm, 15 min, 4 °C) and resuspended in fresh lysis buffer. Cells were lysed using a microfluidizer (18,000 psi, two passes), and the lysate was treated with DNase I followed by two clarifying centrifugations in a Ti-70 rotor (15,000 rpm, 30 min, 4 °C).

The clarified lysate was adjusted to a final concentration of 0.5 M NH₄Cl and layered onto a 1.1 M sucrose cushion prepared in 50 mM Tris-HCl (pH 7.4), 10 mM MgCl₂, 500 mM NH₄Cl, 6 mM β-mercaptoethanol, and 0.5 mM EDTA. Ribosomes were pelleted by ultracentrifugation in a Ti-70 rotor (35,000 rpm, 15 h, 4 °C). The resulting crude ribosome pellet was resuspended in 50 mM Tris-HCl (pH 7.4), 10 mM MgCl₂, 100 mM NH₄Cl, 6 mM β-mercaptoethanol, and 0.5 mM EDTA, followed by ultracentrifugation in a Ti-70 rotor (50,000 rpm, 7 h, 4 °C).

The washed ribosome pellet was resuspended in buffer containing 50 mM Tris-HCl (pH 7.4), 6 mM MgCl₂, 100 mM NH₄Cl, and 6 mM β-mercaptoethanol, and applied to a 10 - 40% (w/v) sucrose gradient prepared in the same buffer. The gradients were centrifuged in a SW-28 rotor (20,000 rpm, 13 hr, 4 °C). Gradients were fractionated using a gradient fractionator equipped with a UA-6 detector (ISCO/BRANDEL). Fractions corresponding to the 70S-MS2-SL peak were pooled, the MgCl₂ concentration was adjusted to 10 mM, and the sample was diluted twofold with 50 mM Tris-HCl (pH 7.4), 10 mM MgCl₂, 100 mM NH₄Cl, and 6 mM β-mercaptoethanol. Ribosomes were pelleted by ultracentrifugation in a Ti-70 rotor (40,000 rpm, 13 h, 4 °C), washed, and resuspended in the same buffer. Ribosome concentration was determined spectrophotometrically using an extinction coefficient of 1 A₂₆₀ = 23 pmol mL⁻¹ for 70S-MS2-SL ribosomes.

### Poly-L-lysine Bead Purification of Ribosome Complexes

A total of 150 pmol *GST–MS2–MNase* protein was incubated with 30 pmol *70S-MS2-SL* ribosomes in binding buffer composed of 50 mM Tris-HCl (pH 7.4), 100 mM NH₄Cl, 10 mM MgCl₂, and 1 mM DTT at room temperature for 30 min. The resulting ribosome complexes were purified using poly-L-lysine magnetic beads. ^38^ Before use, the beads were equilibrated twice with the same binding buffer. The ribosome–protein mixture was incubated with the equilibrated beads for 30 min at 4 °C with gentle agitation.

The beads were collected on a magnetic rack and washed four times with wash buffer containing 50 mM Tris-HCl (pH 7.4), 100 mM NH₄Cl, 10 mM MgCl₂, 1 mM DTT, and 40 µg mL⁻¹ polyglutamic acid. Bound complexes were eluted by incubating the beads in elution buffer composed of 50 mM Tris-HCl (pH 7.4), 100 mM NH₄Cl, 10 mM MgCl₂, 1 mM DTT, and 2 mg mL⁻¹ polyglutamic acid for 15 min at room temperature with shaking. The eluates were analyzed by 10% SDS-PAGE to assess complex formation and used for subsequent MNase cleavage assays or Illumina sequencing library preparation.

### Cleavage Assay of GST-MS2-MNase and Solution MNase with E. coli 70S-MS2-SL Ribosomes

A total of 150 pmol *GST-MS2-MNase* was incubated with 30 pmol *70S-MS2-SL* ribosomes in binding buffer containing 50 mM Tris-HCl (pH 7.4), 100 mM NH₄Cl, 10 mM MgCl₂, and 1 mM DTT. The resulting complex was purified using poly-L-lysine magnetic beads as described above. As a control, a solution MNase reaction was prepared by incubating 30 pmol MNase with 30 pmol *E. coli* 70S ribosomes under identical buffer conditions but without bead purification.

MNase cleavage was initiated by adding CaCl₂ to a final concentration of 1 mM and incubating the reactions at 16 °C for 5 min. Reactions were terminated by the addition of 2 mM EDTA. For negative-control reactions containing inactive MNase, samples were incubated in cleavage buffer (50 mM Tris-HCl, pH 7.4; 100 mM NH₄Cl; 10 mM MgCl₂; 1 mM DTT) at 16 °C for 5 min, followed by quenching with 2 mM EDTA.

Cleaved rRNA fragments were extracted three times with phenol-chloroform (1:1 v/v), followed by two extractions with chloroform alone. The RNA was precipitated with ethanol and resuspended in 20 µL of nuclease-free water. RNA concentration was determined spectrophotometrically using a Tecan Spark plate reader, and the integrity and cleavage pattern were assessed by 6% urea-PAGE. Samples were either analyzed directly or stored for subsequent Illumina sequencing library preparation.

### MNase-FMRP and FMRP-MNase Construct Design and Protein Purification

A plasmid encoding the N-terminally truncated *Drosophila* FMRP (NT-dFMRP, residues 220-681) served as the backbone for fusion construct design. ^36^ Homologous recombination cloning was used to insert the *MNase* gene and linker sequences into the NT-dFMRP plasmid. For the MNase-FMRP construct, the *MNase* coding sequence, followed by a seven-amino acid linker, was inserted between the N-terminal 6×His tag and the NT-dFMRP sequence. For the FMRP-MNase construct, *MNase* was inserted at the C-terminus of NT-dFMRP using a 3×FLAG tag as the linker (22 amino acids).

NT-dFMRP, MNase-FMRP, and FMRP-MNase were expressed and purified using an identical protocol. *E. coli* cells harboring the respective plasmids were cultured overnight in 5 mL LB medium containing 100 µg mL⁻¹ ampicillin at 37 °C. Each 1 L LB culture supplemented with 100 µg mL⁻¹ ampicillin was inoculated with 5 mL of the overnight culture and grown at 37 °C until the optical density at 600 nm reached 0.5. Protein expression was induced with 0.2 mM IPTG, and cultures were incubated for 18 h at 18 °C.

Cells were harvested by centrifugation and resuspended in lysis buffer containing 25 mM Tris-HCl (pH 7.4), 600 mM NaCl, 10 mM imidazole, 10 mM β-mercaptoethanol, and 1 mM PMSF. Lysis was performed by sonication (20 cycles of 8 s at 60% amplitude), and the lysate was clarified by ultracentrifugation in a Ti-70 rotor (22,100 rpm, 30 min, 4 °C). The supernatant was incubated with 3 column volumes of pre-equilibrated Ni–NTA resin for 40 min at 4 °C with gentle mixing. The resin was packed into a gravity column and washed with 15 column volumes of wash buffer composed of 25 mM Tris-HCl (pH 7.4), 600 mM NaCl, 25 mM imidazole, and 4 mM β-mercaptoethanol. Bound proteins were eluted in 10 fractions (1 column volume each) with elution buffer containing 25 mM Tris-HCl (pH 7.4), 600 mM NaCl, 250 mM imidazole, and 10 mM β-mercaptoethanol.

Eluted proteins were further purified by size-exclusion chromatography on a Superdex 200 16/60 column equilibrated in 25 mM Tris-HCl (pH 7.4), 300 mM KCl, and 1 mM DTT. Fractions containing the target protein were identified by SDS-PAGE, pooled, and concentrated using Amicon Ultra centrifugal filters. The final protein samples were stored in 25 mM Tris-HCl (pH 7.4), 300 mM KCl, 1 mM DTT, and 5% (v/v) glycerol at-80 °C. Protein concentrations were determined using the Bradford assay.

### *Drosophila* 80S Ribosome Purification

Dechorionated frozen *Drosophila* embryos were lysed using a Dounce homogenizer in lysis buffer composed of 50 mM HEPES (pH 7.5), 10 mM MgCl₂, 100 mM KCl, 1 mM DTT, 0.5 mM EDTA, 0.5 mM PMSF, 0.1 mM benzamidine, and 200 ng mL⁻¹ heparin. The homogenate was clarified by centrifugation, and the supernatant was layered onto a 30% (w/v) sucrose cushion prepared in the same buffer. Ribosomes were pelleted by ultracentrifugation in a Ti-70 rotor at 39,000 rpm for 16 h at 4 °C.

The crude ribosome pellet was resuspended in buffer containing 50 mM HEPES (pH 7.5), 2 mM MgCl₂, 500 mM KCl, 1 mM DTT, 0.5 mM PMSF, 0.1 mM benzamidine, and 200 ng mL⁻¹ heparin, followed by centrifugation in a Ti-70 rotor at 13,000 rpm for 20 min at 4 °C to remove insoluble material. To release nascent polypeptides, puromycin was added to the supernatant at a ratio of 1 mg puromycin per 100 mg ribosome, and the mixture was incubated for 1 h on ice, followed by 25 min at 37 °C.

The lysate was layered onto a 10 - 40% (w/v) sucrose gradient and centrifuged in a SW-28 rotor (20,700 rpm, 15 hr, 4 °C), and fractionated using a gradient fractionator equipped with a UA-6 detector (ISCO/BRANDEL). Fractions corresponding to the 80S ribosome peak were pooled, diluted twofold with buffer containing 50 mM HEPES (pH 7.5), 2 mM MgCl₂, 25 mM KCl, and 1 mM DTT, and centrifuged in a Ti-70 rotor at 27,500 rpm for 19 h at 4 °C. The purified ribosome pellet was resuspended in storage buffer composed of 10 mM HEPES (pH 7.5), 10 mM MgCl₂, and 50 mM KCl, flash-frozen, and stored at-80 °C. Ribosome concentration was determined spectrophotometrically using an extinction coefficient of 1 A₂₆₀ unit = 20 pmol mL⁻¹ for 80S ribosomes.

### Electrophoretic Mobility Shift Assay (EMSA) of NT-dFMRP, MNase-FMRP, and FMRP-MNase with PolyG₁₈ and CR1

Binding reactions were performed by titrating NT-dFMRP, MNase-FMRP, or FMRP-MNase against 100 nM fluorescein-labeled PolyG₁₈ or CR1 RNA. Each 26 µL reaction contained 50 mM Tris-HCl (pH 7.4), 130 mM KCl, 5 mM MgCl₂, 25 µg mL⁻¹ BSA, 10 mM DTT, and 50 µg mL⁻¹ tRNA as a nonspecific competitor. Mixtures were incubated at room temperature for 1 h in the dark, followed by the addition of 3 µL of 50% (v/v) glycerol containing xylene cyanol tracking dye.

Samples were resolved on 0.8% (w/v) SeaKem GTG agarose gels prepared in 1× TBE buffer and electrophoresed at 66 V for 1 h at 4 °C. Fluorescent bands were visualized using a Typhoon FLA 9500 scanner (Cytiva), and band intensities were quantified using *ImageJ* software (National Institutes of Health).

### Fluorescence Anisotropy Binding Assay and Determination of Equilibrium Dissociation Constants for NT-dFMRP, MNase-FMRP, and FMRP-MNase

Fluorescence anisotropy measurements were used to determine the equilibrium dissociation constants (*K*_D_) of NT-dFMRP, MNase-FMRP, and FMRP-MNase binding to fluorescein-labeled PolyG₁₈ and CR1 RNAs. Protein samples were titrated against 10 nM of fluorescein-labeled RNA in a total reaction volume of 100 µL. Binding reactions were performed in buffer containing 50 mM Tris-HCl (pH 7.4), 130 mM KCl, 5 mM MgCl₂, 25 µg mL⁻¹ BSA, 10 mM DTT, and 50 µg mL⁻¹ tRNA. The mixtures were incubated at room temperature for 1 h in the dark to reach equilibrium.

Fluorescence anisotropy was measured in non-binding, black, flat-bottom 96-well plates (Greiner) using a Tecan Spark multimode microplate reader. Samples were excited at 485 nm (20 nm bandwidth), and emission was recorded at 535 nm (25 nm bandwidth). Optimal gain and Z-position parameters were determined automatically for each well.

The change in anisotropy was plotted as a function of protein concentration, and the data were fit to the quadratic equation below using GraphPad Prism (GraphPad Software, San Diego, CA) to obtain the equilibrium dissociation constant (*K*_D_).

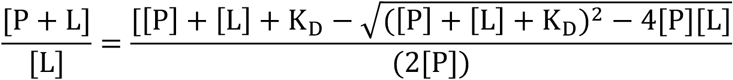

Here [P+L]/[L] represents the anisotropy value, [L] is the ligand concentration, and [P] is the protein concentration. The *K*_D_ value was determined as the average of three independent trials.

### Binding Assay of NT-dFMRP, MNase-FMRP, and FMRP-MNase with *Drosophila* 80S Ribosomes

Binding reactions were performed by incubating 125 pmol of MNase-FMRP or FMRP-MNase with 25 pmol of *Drosophila* 80S ribosomes in binding buffer composed of 50 mM Tris-HCl (pH 7.4), 5 mM MgCl₂, 1 mM DTT, and 50 µg mL⁻¹ tRNA. Reactions were carried out at room temperature for 30 min.

Ribosome-protein complexes were purified by ultracentrifugation through a 1.1 M sucrose cushion prepared in 50 mM Tris-HCl (pH 7.4), 50 mM KCl, 5 mM MgCl₂, and 1 mM DTT. Each 20 µL reaction was layered onto 200 µL of sucrose cushion and centrifuged in a Ti-42.2 rotor at 38,000 rpm for 1 h 40 min at 4 °C. Pelleted complexes were gently resuspended in buffer containing 50 mM Tris-HCl (pH 7.4), 50 mM KCl, 5 mM MgCl₂, and 1 mM DTT.

Samples were analyzed by 10% SDS-PAGE to assess ribosome binding and subsequently used for MNase cleavage assays or Illumina sequencing library preparation.

### Cleavage Assay of MNase-FMRP, FMRP-MNase, and Solution MNase with *Drosophila* 80S Ribosomes

A total of 125 pmol MNase–FMRP or FMRP–MNase was incubated with 25 pmol *Drosophila* 80S ribosomes in binding buffer containing 50 mM Tris-HCl (pH 7.4), 50 mM KCl, 5 mM MgCl₂, and 1 mM DTT. The resulting ribosome complexes were purified by ultracentrifugation through a sucrose cushion as described above. For control reactions with free (solution) MNase, 25 pmol MNase and 25 pmol *Drosophila* 80S ribosomes were incubated under identical buffer conditions without sucrose cushion purification.

MNase activity was initiated by adding CaCl₂ to a final concentration of 1 mM and incubating the reactions at 16 °C for 5 min. Reactions were terminated by the addition of 2 mM EDTA. For inactive MNase controls, samples were incubated under the same buffer conditions at 16 °C for 5 min without CaCl₂, followed by quenching with 2 mM EDTA.

Cleaved rRNA fragments were extracted three times with phenol-chloroform (1:1, v/v) and twice with chloroform. RNA was precipitated with ethanol and resuspended in 20 µL of nuclease-free water. rRNA concentrations were determined using a Tecan Spark plate reader, and cleavage patterns were analyzed by 6% urea-PAGE. Purified rRNA was either analyzed directly or stored for subsequent Illumina sequencing library preparation.

### Illumina Sequencing Library Preparation

A total of 5 µg of purified rRNA was used for dephosphorylation in a 50 µL reaction containing 5 µL CutSmart Buffer (New England Biolabs), 2 µL recombinant shrimp alkaline phosphatase (rSAP), and 1 µL murine RNase inhibitor. Reactions were incubated at 37 °C for 30 min with gentle agitation, and enzymes were heat-inactivated at 65 °C for 5 min. RNA was purified using RNAClean XP beads (Beckman Coulter) and eluted in 16 µL of nuclease-free water. RNA concentration was determined spectrophotometrically using a Tecan Spark plate reader.

For poly(A) tailing, 1 µg of dephosphorylated rRNA was incubated in a 20 µL reaction containing 2 µL E. coli Poly(A) Polymerase Reaction Buffer, 2 µL 10 mM ATP, and 1 µL E. coli poly(A) polymerase (New England Biolabs). Reactions were incubated at 37 °C for 20 min and terminated by adding EDTA to a final concentration of 12.5 mM. RNA was purified with RNAClean XP beads and eluted in 8 µL of nuclease-free water.

Polyadenylated RNA was fragmented by the addition of 1 µL 10× RNA fragmentation reagent (Ambion), incubation at 70 °C for 5 min, and immediate quenching with 1 µL stop solution.

Approximately 200 ng of the fragmented, poly(A)-tailed RNA was annealed with the Oligo(dT)₃₀VN_REV primer (5’-GTCTCGTGGGCTCGGAGATGTGTATAAGAGACAGT_30_VN-3’) at 65 °C, followed by addition of the template-switching primer (5’-TCGTCGGCAGCGTCAGATGTGTATAAGAGACAG**rGrGrG**-3’), template-switching reverse-transcription buffer, and enzyme mix (Catalog# M0466S, New England Biolabs). Reverse transcription was carried out at 42 °C for 90 min and terminated at 85 °C for 5 min. The resulting cDNA was treated with 1 µg RNase A for 30 min at room temperature and purified with 1× RNAClean XP beads, followed by two washes with 80% ethanol.

Purified cDNA was eluted in nuclease-free water and amplified using Illumina i5 and i7 index primers with KOD Hot Start DNA polymerase (Millipore Sigma). PCR products were purified using 0.8× RNAClean XP beads with two 80% ethanol washes, and the final libraries were eluted in 17 µL Illumina Resuspension Buffer (RSB). Library concentrations were quantified using the Qubit dsDNA HS Assay Kit (Thermo Fisher Scientific). Libraries were pooled equimolarly, multiplexed, and sequenced on an Illumina NovaSeq X Plus 10B platform, generating on average 30 million paired-end reads per sample.

### Illumina Sequencing Data Processing

Raw Illumina sequencing reads were first screened for sequences containing ≥30 consecutive adenosine (A) residues, corresponding to poly(A) tails. Reads meeting this criterion were retained, and poly(A) tails were trimmed before alignment. The filtered and trimmed reads were then aligned separately to the *E. coli* 23S and 16S rRNA reference sequences using Salmon (version 1.10.3) with default parameters. ^39^

Following alignment, the nucleotide position of the 3′ end of each mapped read was extracted using custom Python scripts developed to identify and count unique cleavage positions across the rRNA sequence. The resulting 3′-end counts were normalized relative to the counts at the authentic 3′ termini of the respective rRNAs to correct for sequencing depth and sample variation. The same analytical workflow was applied to identify MNase-dependent cleavage sites within the *Drosophila* 28S and 18S rRNAs. The cleavage sites were then mapped onto high-resolution ribosome structures using UCSF ChimeraX. ^40^

## RESULTS

### Mapping RBP-RNA interactions using an MNase-MS2 fusion strategy

We engineered an MS2-MNase fusion protein to develop a proximity-based method for mapping the location of RNA-binding protein (RBP) on RNA at nucleotide resolution. The MS2 coat protein recognizes and binds its cognate RNA stem-loop with exceptionally high affinity, a feature that has previously been exploited to purify *E. coli* ribosomes by inserting MS2 stem-loops into the 23S or 16S rRNA and capturing the tagged subunits on MS2-GST affinity matrices. ^27^ Building on this concept, we selected *E. coli* ribosomes containing an MS2 stem-loop engineered into helix 98 of the 23S rRNA to validate our approach. ^28^ This position was chosen because a high-resolution structure of the ribosome with the MS2 stem-loop is available, providing a well-defined spatial framework for assessing cleavage specificity. ^30^ In our experimental design, the MS2-MNase fusion binds to the MS2 stem-loop on the ribosome to form a ribosome•MS2-MNase complex. Upon activation of MNase by Ca²⁺, RNA regions located within the accessible radius of MNase are selectively cleaved. The resulting rRNA fragments can then be extracted, sequenced, and mapped to reveal MNase-dependent cleavage sites and thereby delineate the local RNA-protein interaction landscape (Figure 1A).

**Figure 1.**
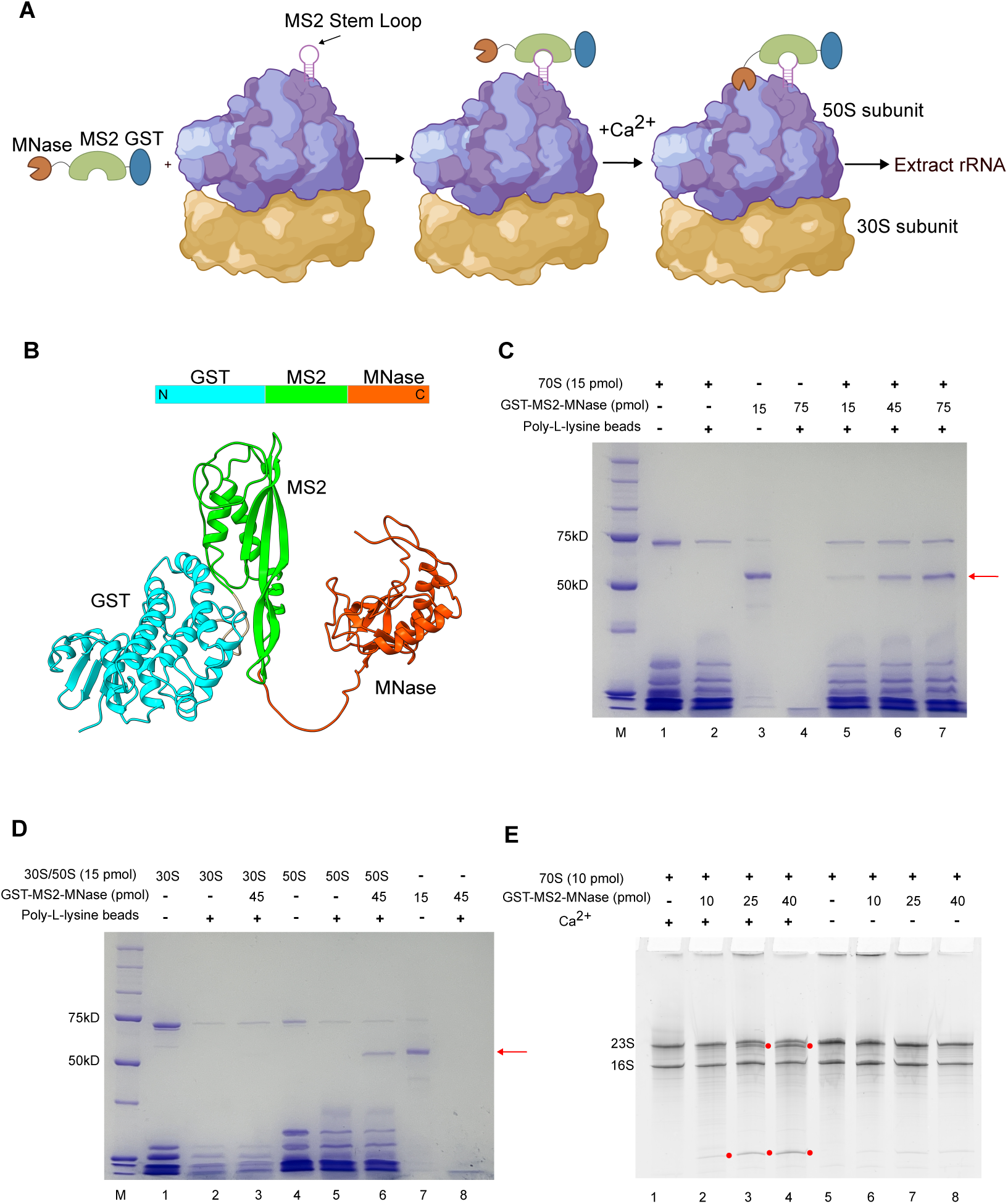
Design and characterization of the GST-MS2-MNase fusion protein and its interaction with the E. coli ribosome. **(A)** Schematic representation of the tethered MNase mapping strategy. The MS2-MNase fusion binds specifically to an engineered MS2 stem-loop (MS2-SL) inserted on the surface of the 50S ribosomal subunit. Upon addition of Ca²⁺, MNase becomes activated and cleaves nearby rRNA regions. The resulting rRNA fragments are extracted and sequenced to identify MNase-dependent cleavage sites at nucleotide resolution. Created using BioRender. **(B)** Predicted three-dimensional structure of GST-MS2-MNase generated by AlphaFold2. The model shows the GST (cyan), MS2 (green), and MNase (orange) domains connected by flexible linkers, consistent with the modular architecture of the fusion protein. **(C)** Binding of GST-MS2-MNase to 70S ribosomes containing the engineered MS2-SL. Complexes were purified using poly-L-lysine beads and analyzed by SDS-PAGE. Lanes: M, molecular-weight marker; 1, input 70S ribosome; 2, 70S ribosome recovered after purification; 3, GST-MS2-MNase control; 4, GST-MS2-MNase removed by purification; 5 - 7, 70S ribosomes incubated with increasing amounts of GST-MS2-MNase and purified. The red arrow marks GST-MS2-MNase. **(D)** Binding specificity of GST-MS2-MNase to isolated 30S and 50S subunits. Lanes: M, molecular-weight marker; 1 - 3, 30S controls and incubations; 4 - 6, 50S controls and incubations; 7 - 8, GST-MS2-MNase controls. The red arrow marks GST-MS2-MNase. **(E)** Cleavage of 23S rRNA by GST-MS2-MNase bound to 70S ribosomes. The 70S-MS2-SL•GST-MS2-MNase complex was purified, activated with Ca²⁺, and analyzed by denaturing PAGE. Lanes: 1 - 8 as indicated; red circles denote MNase-dependent 23S rRNA cleavage products.

To generate a functional construct, we designed a tripartite fusion protein comprising glutathione S-transferase (GST), MS2 coat protein, and MNase (GST-MS2-MNase). Structural prediction using AlphaFold2 suggested that GST, MS2, and MNase each fold as independent, stable domains connected by flexible linkers (Figure 1B). ^41^ The model further indicated that the six-amino acid linker between MS2 and MNase provides sufficient conformational flexibility for MNase to sample RNA regions within roughly 20 Å of the MS2 RNA stem-loop. This architecture was expected to permit localized cleavage while maintaining specific tethering of MNase to the ribosome through the MS2 stem-loop interaction.

We next examined whether the GST-MS2-MNase fusion protein associates specifically with *E. coli* 70S ribosomes carrying the engineered MS2 stem-loop (70S-MS2-SL). To test this, 70S-MS2-SL complexes were incubated with increasing concentrations of GST-MS2-MNase and purified using poly-L-lysine beads, which selectively capture ribosomes and enable removal of unbound proteins. ^38^ The bead-bound material was analyzed by SDS-PAGE. As shown in Figure 1C, 70S ribosomes efficiently bound and eluted from the beads (lane 2), whereas GST-MS2-MNase alone did not (lane 4). Strikingly, when 70S-MS2-SL was incubated with GST-MS2-MNase, the fusion protein co-eluted with the ribosomes in a concentration-dependent manner (lanes 5-7), indicating stable complex formation between the ribosome and the fusion protein.

These results confirm that GST-MS2-MNase associates with ribosomes containing the engineered MS2 stem-loop.

To further validate the specificity of this interaction, we analyzed the binding of GST-MS2-MNase to the individual ribosomal subunits. The 50S subunit, which contains the MS2 stem-loop insertion, and the 30S subunit, which lacks it, were each incubated with GST-MS2-MNase, purified on poly-L-lysine beads, and analyzed by SDS-PAGE. Consistent with specific recognition of the MS2 stem-loop, GST-MS2-MNase bound robustly to the 50S subunit but not to the 30S subunit (Figure 1D, lanes 3 and 6). Together, these results demonstrate that GST-MS2-MNase binds selectively to the MS2-tagged 50S subunit within the assembled 70S ribosome.

To determine whether the MNase domain retained catalytic activity in the context of the ribosome-bound complex, we incubated the poly-L-lysine bead-purified 70S-MS2-SL•GST-MS2-MNase assembly with Ca²⁺ to activate MNase and analyzed the resulting rRNAs by denaturing polyacrylamide gel electrophoresis. Distinct cleavage fragments were observed only in reactions containing Ca²⁺ (Figure 1E, lanes 2 - 4), demonstrating that MNase remained enzymatically competent when tethered to the ribosome. No cleavage products were detected in the absence of Ca²⁺, confirming the reaction’s dependence on MNase activation. Importantly, cleavage occurred in the 23S rRNA but not in the 16S rRNA, consistent with the localization of the MS2 stem-loop insertion and the binding site of GST-MS2-MNase within the 23S rRNA.

### Mapping MS2-MNase cleavage sites on rRNA by high-throughput sequencing

To precisely determine the sites of MNase cleavage directed by the tethered fusion protein, we established a high-throughput sequencing workflow. Following the Ca²⁺-dependent cleavage reaction, rRNAs were purified, dephosphorylated with alkaline phosphatase, and polyadenylated using poly(A) polymerase. The addition of the poly(A) tail served as a molecular tag marking the 3′ ends of MNase-generated fragments. To construct Illumina sequencing libraries, the rRNA was further fragmented non-specifically to ∼60–200 nucleotides using RNA Fragmentation Reagent. The resulting fragments were reverse-transcribed with an oligo(dT)₃₀ primer and a template-switching primer, both of which contained Illumina adapter sequences. The cDNA was then amplified using Illumina index primers to generate barcoded libraries for sequencing.

Sequencing reads were processed to remove low-quality reads, trimmed of their poly(A) tails, and aligned to the *E. coli* 16S and 23S rRNA reference sequences. The 3′ termini of reads were extracted using a custom Python script and plotted along the rRNA sequence (Figure 2A, 2B, and 2C). As expected, the majority of 3′ ends mapped to the authentic 3′ termini of the 16S and 23S rRNAs (Figure 2A and 2B, black bar). Only background-level cleavage was observed in the 16S rRNA. Remarkably, in the sample containing the 70S-MS2-SL•GST-MS2-MNase complex and Ca²⁺, we detected a prominent accumulation of 3′ ends at position U545 of the 23S rRNA (Figure 2B and 2C, red bar). In contrast, this cleavage peak was absent in the control lacking Ca²⁺, as well as in the reaction where free, untethered MNase was added to the 70S-MS2-SL ribosomes. These findings demonstrate that GST-MS2-MNase specifically associates with the MS2 stem-loop inserted in the 70S ribosome and catalyzes a localized cleavage at position U545 of the 23S rRNA. The absence of signal in the Ca²⁺-free and untethered MNase controls confirms that this cleavage is both Ca²⁺-dependent and dependent on the targeted recruitment of MNase via the MS2-RNA interaction.

**Figure 2.**
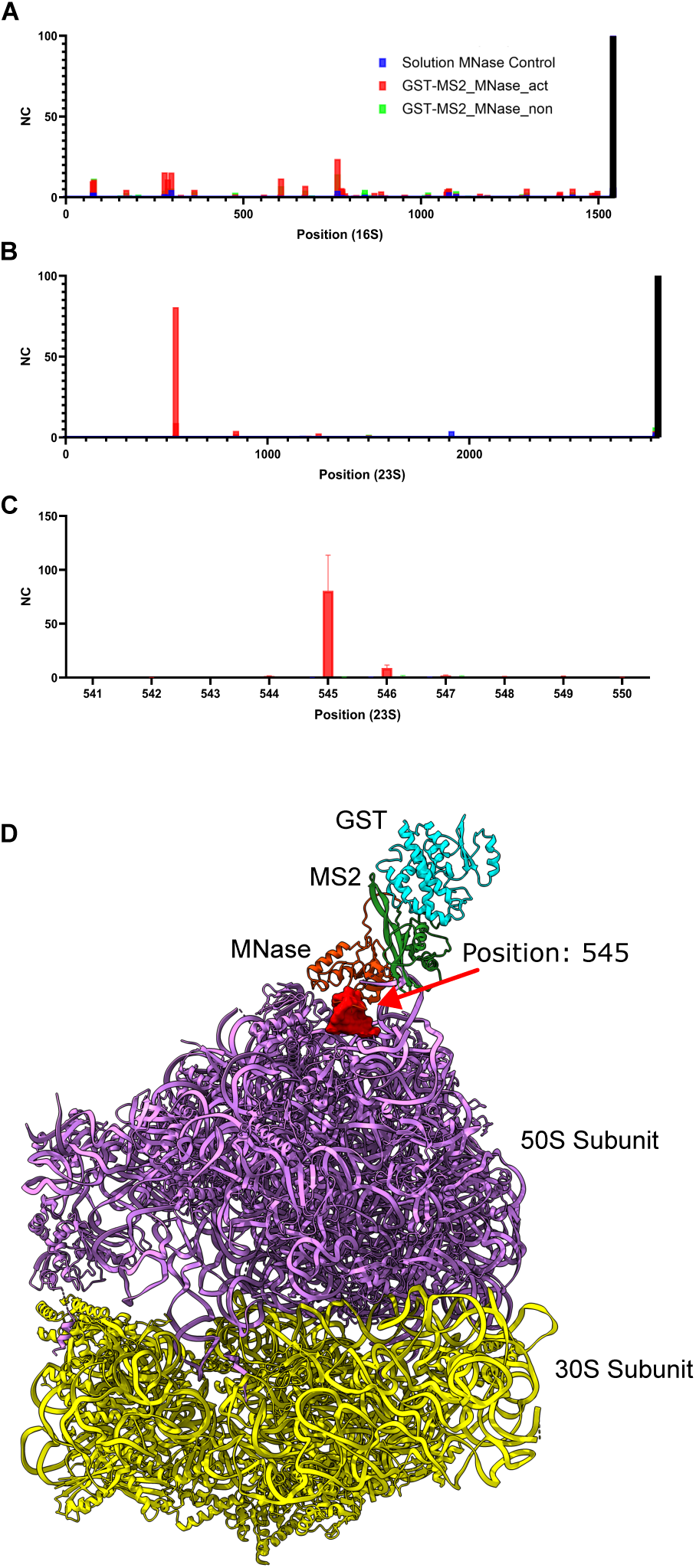
Mapping of MS2-MNase cleavage sites on rRNA by high-throughput sequencing. **(A)** Distribution of normalized 3′-termini read counts mapped to the *E. coli* 16S rRNA sequence. The position of each bar corresponds to a nucleotide in the rRNA, and the height reflects the normalized number of reads ending at that position. The error bars indicate the SD from three independent experiments. **(B)** Distribution of normalized 3′-termini read counts mapped to the 23S rRNA sequence. The prominent red bar indicates the site of MNase-dependent cleavage near the engineered MS2 stem-loop. The error bars indicate the SD from three independent experiments. **(C)** Close-up view of the region surrounding nucleotide 545 of the 23S rRNA, where strong MNase-dependent cleavage was observed. The x-axis denotes the nucleotide position, and the y-axis shows normalized counts (NC). Read counts were normalized by setting the total counts for the authentic 3′-termini of the rRNAs to 100 (black bar). Cleavage patterns are shown for soluble MNase control (blue bars), GST-MS2-MNase + Ca²⁺ (red bars), and GST-MS2-MNase without Ca²⁺ (green bars). **(D)** Structural localization of MNase cleavage sites on the *E. coli* 70S-MS2-SL•GST-MS2-MNase complex. The modeled GST-MS2-MNase fusion was positioned on helix 98 of the 23S rRNA, and the primary cleavage site at position 545 is highlighted in red. The structural mapping confirms that MNase cleaves rRNA regions spatially proximal to the MS2-SL insertion site, validating the tethered nuclease mapping approach. (70S-MS-SL structure PDB ID: 8FTO)

Having identified a prominent MNase-dependent cleavage site near the engineered MS2 stem-loop, we next examined its spatial position on the ribosome to assess the accuracy of the tethered mapping approach. Mapping the cleavage site at position U545 onto the *E. coli* ribosome structure revealed that it lies within approximately ∼15 Å of the MS2 stem-loop, confirming that the MNase-MS2 fusion faithfully reports the three-dimensional location of the bound RNA-binding protein (Figure 2D). Interestingly, the cleavage was localized to one side of the MS2 stem-loop, suggesting that the attached GST domain restricts the rotational freedom of the MNase, thereby limiting its access to surrounding rRNA regions. Alternatively, this asymmetric cleavage pattern may arise from the absence of other accessible rRNA elements in the immediate vicinity of the MNase active site.

### MNase Fusion Design and RNA-Binding Characterization of *Drosophila* FMRP

Building on the successful validation of the TM-map approach using the MS2-MNase fusion, we next applied the method to define the ribosome-binding site of *Drosophila* FMRP on the 80S ribosome. Previous biochemical studies have shown that FMRP associates with ribosomes and acts as a translational repressor. ^31–36^ Moreover, a cryo-EM structure of the N-terminal truncated form of *Drosophila* FMRP (NT-dFMRP) bound to the ribosome revealed density in the intersubunit space, beneath the central protuberance of the 60S subunit. ^36^ However, the limited resolution of that structure precluded precise localization of the protein on the ribosome surface. To refine the positional information, we engineered two MNase fusion constructs, in which MNase was attached either to the N-or C-terminus of NT-dFMRP (Figure 3A). We used AlphaFold2 to predict the three-dimensional structure of each fusion and assess the relative spatial placement of the MNase domain (Figure 3B and 3C). ^41^ The models suggested that MNase fused to the N-terminus, adjacent to the structured KH1 and KH2 domains, may have restricted mobility and limited sampling of the surrounding ribosomal environment. In contrast, the C-terminally fused MNase is likely to possess greater flexibility, as the C-terminal region of NT-dFMRP is largely disordered, allowing broader access to nearby rRNA elements.

**Figure 3.**
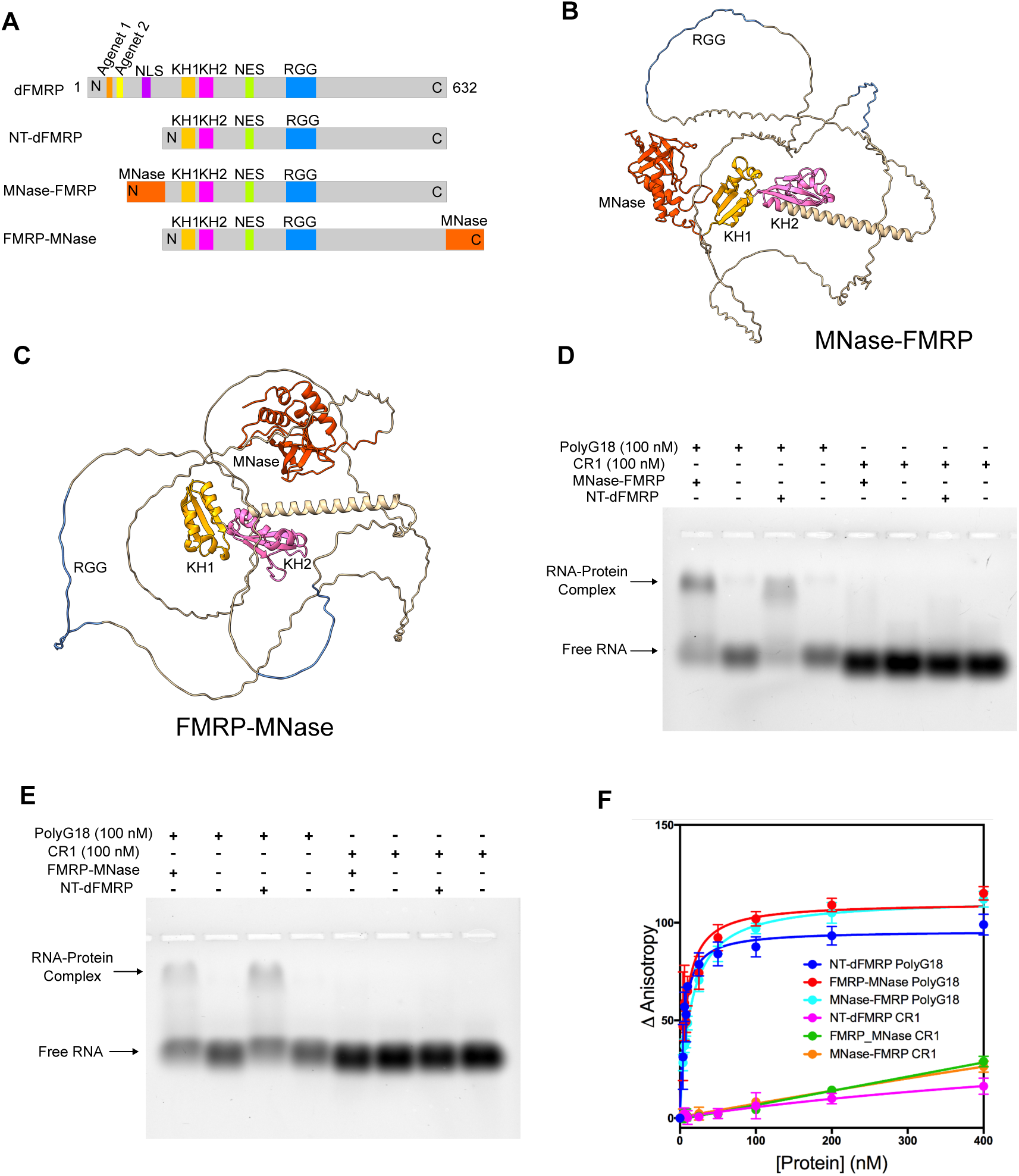
Design and characterization of MNase-FMRP and FMRP-MNase fusion proteins. **(A)** Schematic representation of the domain organization in *Drosophila* FMRP (dFMRP), the N-terminal truncated FMRP (NT-dFMRP), and the MNase-FMRP and FMRP-MNase fusion proteins. Functional domains include the Agenet 1 and 2 domains, K-homology (KH) RNA-binding domains (KH1 and KH2), and the arginine-glycine-glycine (RGG) motif within the disordered C-terminal region. **(B)** Predicted three-dimensional structure of MNase-FMRP generated by AlphaFold2. The model shows the N-terminal MNase domain (orange) fused to FMRP, highlighting KH1 (yellow), KH2 (magenta), and the RGG motif in the disordered region. **(C)** Predicted structure of FMRP-MNase generated by AlphaFold2. The model shows FMRP with KH1 (yellow), KH2 (magenta), and the RGG motif in the disordered region, followed by the C-terminal MNase domain (orange). **(D)** Electrophoretic mobility shift assay (EMSA) showing that MNase-FMRP binds specifically to polyG₁₈ RNA but not to the CR1 control RNA. The presence (+) or absence (–) of RNA and protein is indicated above each lane. The arrows mark the positions of free RNA and RNA-protein complex. **(E)** EMSA analysis of FMRP-MNase showing binding to polyG₁₈ RNA but not to CR1 RNA, as indicated by the shifted RNA-protein complex. **(F)** Fluorescence anisotropy analysis of RNA binding. Increasing concentrations of NT-dFMRP, MNase-FMRP, or FMRP-MNase were titrated against fluorescently labeled polyG₁₈ RNA or CR1 RNA. The x-axis shows protein concentration, and the y-axis shows anisotropy change (arbitrary units). All three proteins bound polyG₁₈ RNA with comparable apparent *K*D values but did not bind CR1 RNA.

The recombinant MNase-FMRP and FMRP-MNase fusion proteins were expressed and purified from *E. coli*. To confirm that the addition of the MNase domain did not interfere with the intrinsic RNA-binding properties of NT-dFMRP, we compared the RNA-binding activity of the fusion proteins with that of the wild-type NT-dFMRP. This verification was essential to ensure that the MNase fusions remained functionally competent for downstream TM-map analysis. We assessed RNA binding using an electrophoretic mobility shift assay (EMSA) with two previously characterized RNA substrates: polyG₁₈, which forms a G-quadruplex (GQ) structure recognized by NT-dFMRP, and CR1, a non-binding control RNA. ^42^ In the presence of polyG₁₈, all three proteins: NT-dFMRP, MNase-FMRP, and FMRP-MNase produced a slower-migrating band corresponding to the RNA-protein complex (Figure 3D and 3E). No shift was observed with CR1, confirming the specificity of the interaction. Together, these results demonstrate that both MNase fusion proteins retain RNA-binding activity comparable to that of NT-dFMRP, indicating that attachment of MNase does not disrupt FMRP’s native RNA recognition properties.

Having established that the MNase fusions retain RNA-binding activity, we next quantified their binding affinities relative to wild-type NT-dFMRP. Fluorescence anisotropy assays were performed using fluorescein-labeled polyG₁₈ and CR1 RNAs. ^42^ Increasing concentrations of NT-dFMRP, MNase-FMRP, or FMRP-MNase were titrated against the labeled RNAs, and the resulting changes in fluorescence anisotropy were used to calculate the equilibrium dissociation constants (*K*D). The binding curves showed a characteristic increase in anisotropy with rising protein concentrations for polyG₁₈, but not for the non-binding CR1 RNA (Figure 3F). The calculated *K*D values for NT-dFMRP, MNase-FMRP, and FMRP-MNase were 5 ± 1 nM, 14 ± 1 nM, and 8 ± 2 nM, respectively. These data indicate that both fusion proteins bind polyG₁₈ with high affinity comparable to that of NT-dFMRP. Although the attachment of MNase modestly reduced binding strength, the effect was minor, confirming that the MNase fusions remain functionally competent for RNA recognition and suitable for subsequent ribosome-mapping experiments.

### Ribosome Association and Catalytic Activity of MNase-FMRP Fusion Proteins

Previous studies have shown that NT-dFMRP binds directly to *Drosophila* 80S ribosomes even in the absence of mRNA, as demonstrated by sucrose cushion centrifugation assays. ^36^ To confirm that the MNase-FMRP and FMRP-MNase fusion proteins retain this ribosome-binding ability, we performed parallel ultracentrifugation experiments using purified 80S ribosomes (Figure 4A). Ribosome-FMRP complexes were assembled in vitro, layered onto sucrose cushions, and subjected to ultracentrifugation to separate ribosome-bound from unbound protein. Proteins that associate with the ribosome are expected to co-sediment into the pellet, whereas unbound proteins remain in the supernatant. Analysis of the resuspended ribosome pellets by SDS-PAGE revealed distinct bands corresponding to MNase-FMRP and FMRP-MNase fusion proteins (71 kDa) (Figure 4B). These results demonstrate that both MNase fusions stably associate with the 80S ribosome and that the ribosome-FMRP complexes can be efficiently recovered by sucrose cushion centrifugation.

**Figure 4.**
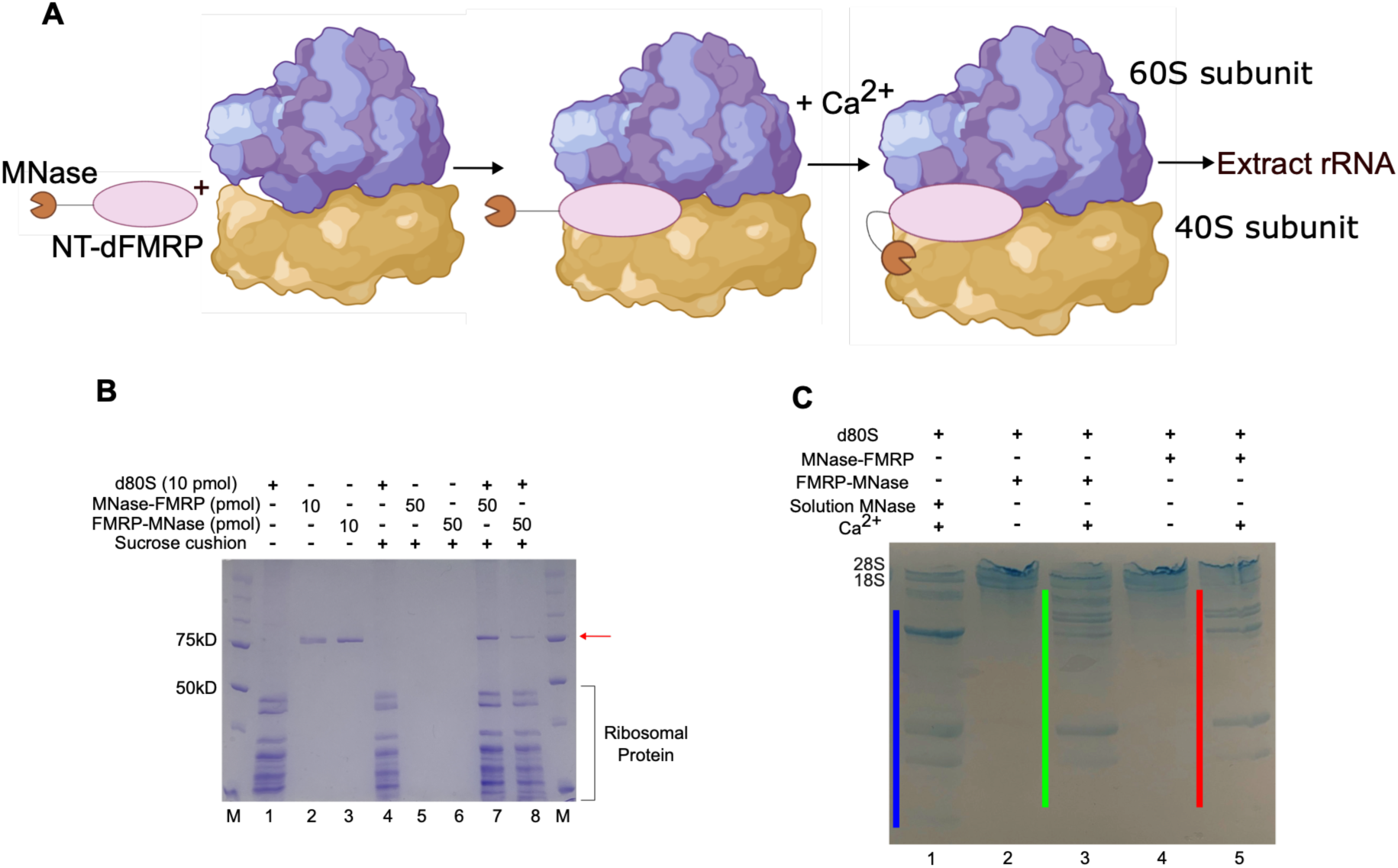
Interaction of MNase-FMRP and FMRP-MNase fusion proteins with the *Drosophila* 80S ribosome. **(A)** Schematic representation of the tethered MNase mapping strategy applied to the *Drosophila* 80S ribosome. The MNase-FMRP fusion binds to the ribosome, and upon addition of Ca²⁺, MNase becomes activated and cleaves proximal rRNA regions. The resulting rRNA fragments are extracted and sequenced to identify MNase-dependent cleavage sites at nucleotide resolution. Created using BioRender. **(B)** Analysis of MNase-FMRP and FMRP-MNase binding to *Drosophila* 80S ribosomes (d80S). Complexes were purified through sucrose cushion ultracentrifugation and analyzed by SDS-PAGE. Lanes: M, molecular-weight marker; 1, input d80S ribosome; 2, input MNase-FMRP; 3, input FMRP-MNase; 4, purified d80S ribosome; 5, MNase-FMRP removed by purification; 6, FMRP-MNase removed by purification; 7, d80S ribosomes incubated with MNase-FMRP and purified; 8, d80S ribosomes incubated with FMRP-MNase and purified. The red arrow indicates the position of the MNase-FMRP and FMRP-MNase fusion proteins. **(C)** Ca²⁺-dependent cleavage of rRNAs by MNase-FMRP and FMRP-MNase bound to d80S ribosomes. The d80S•MNase-FMRP and d80S•FMRP-MNase complexes were purified using sucrose cushions, activated with Ca²⁺, and analyzed by denaturing PAGE. Lanes 1 - 5 correspond to the conditions indicated in the figure. The blue line represents cleavage by soluble MNase, the green line indicates cleavage by FMRP-MNase, and the red line indicates cleavage by MNase-FMRP. Distinct cleavage patterns confirm that both fusion proteins associate with ribosomes and direct MNase activity toward neighboring rRNA regions.

Having verified that MNase-FMRP and FMRP-MNase retained both RNA-and ribosome-binding activity, we next examined whether the MNase fusion proteins were catalytically active when bound to the ribosome. Ribosome-FMRP complexes were purified by sucrose cushion ultracentrifugation, resuspended in buffer, and divided into aliquots. MNase activity was initiated by adding Ca²⁺ to one set of each complex, while parallel samples without Ca²⁺ served as negative controls to assess background RNA cleavage. To control for nonspecific digestion by unbound enzyme, we also incubated purified ribosomes with soluble MNase, which does not associate with the ribosome. Cleavage reactions were carried out for 5 min at 16 °C and terminated by adding excess EDTA. The resulting rRNAs were purified and analyzed by denaturing PAGE. As shown in Figure 4C, Ca²⁺-activated MNase-FMRP (red bar) and FMRP-MNase (green bar) samples displayed multiple distinct rRNA fragments smaller than the intact rRNAs, whereas control samples lacking Ca²⁺ showed little or no cleavage. Soluble MNase also produced cleavage fragments (blue bar), but the banding pattern differed markedly from that of the tethered fusions. These results demonstrate that the MNase-FMRP and FMRP-MNase fusion proteins are catalytically active and generate ribosome-associated rRNA cleavages consistent with the site-specific activity of the tethered MNase.

### TM-Map Reveals MNase-Dependent Cleavage Patterns of Ribosome-Bound FMRP

Having confirmed the activity of the MNase-FMRP fusions, we next sought to map the precise locations of these cleavage events on the ribosome. To do this, we extracted the cleaved rRNAs from each reaction, generated poly(A)-tailed sequencing libraries, and performed high-throughput RNA sequencing. The sequencing data were processed to identify the 3′ ends of the cleaved fragments, allowing us to pinpoint MNase-dependent cleavage sites at nucleotide resolution and visualize their distribution across the *Drosophila* rRNAs (Figure 5).

**Figure 5.**
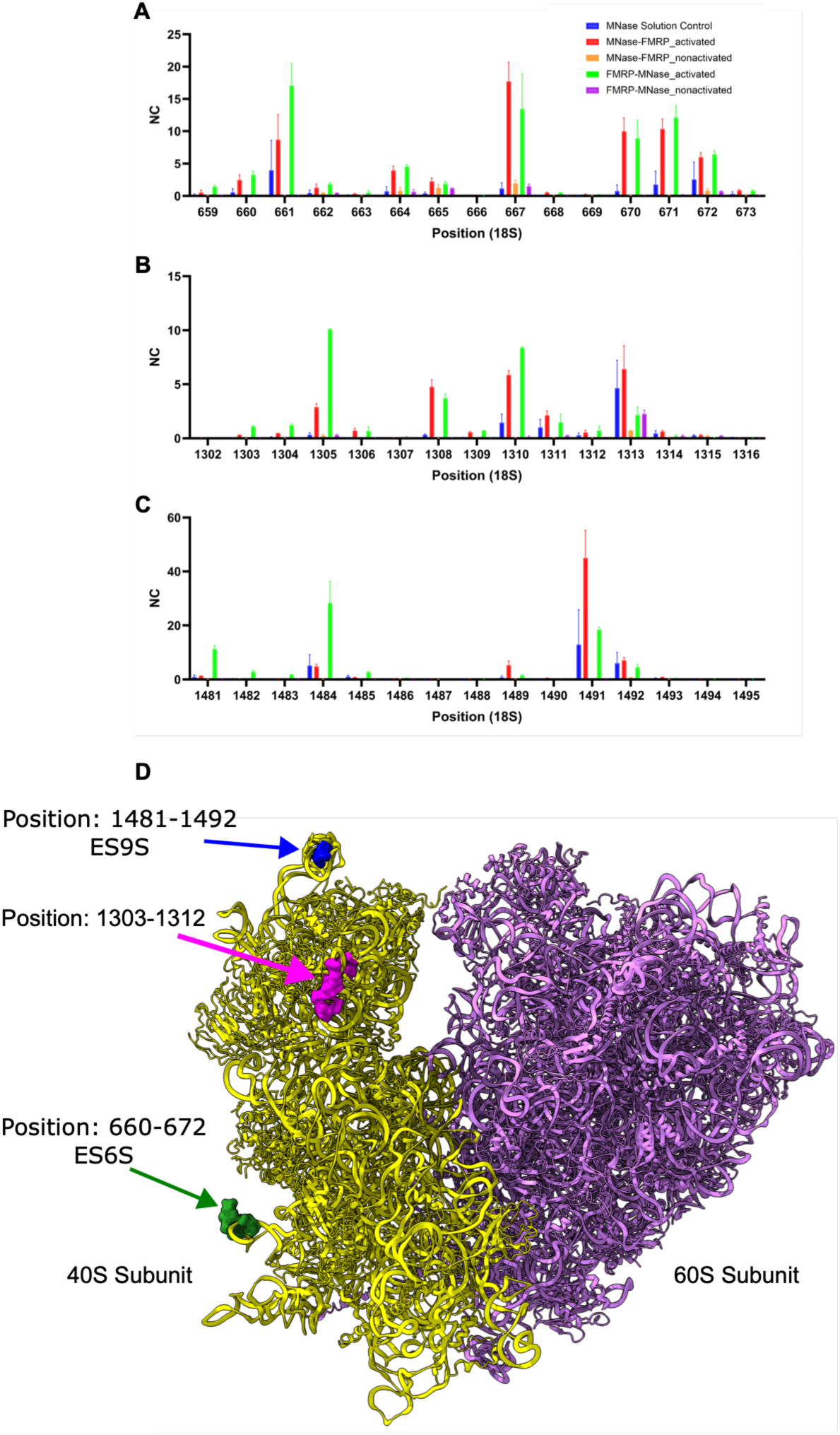
Mapping of MNase-FMRP and FMRP-MNase cleavage sites on *Drosophila* rRNA by high-throughput sequencing. **(A - C)** Distribution of normalized 3′-termini read counts mapped to selected regions of the *Drosophila* 18S rRNA. Panels show cleavage profiles spanning nucleotides 659 - 673 (A), 1302 - 1316 (B), and 1481 - 1495 (C). The x-axis indicates nucleotide position along the rRNA, and the y-axis represents normalized counts (NC) corresponding to the number of sequencing reads terminating at each position. Read counts were normalized by setting the total for authentic 3′ termini to 100. Cleavage patterns are shown for soluble MNase control (blue bars), MNase-FMRP + Ca²⁺ (red bars), MNase-FMRP without Ca²⁺ (orange bars), FMRP-MNase + Ca²⁺ (green bars), and FMRP-MNase without Ca²⁺ (purple bars). The error bars indicate the SD from two independent experiments. Distinct clusters of MNase-dependent cleavage were observed in the MNase-FMRP and FMRP-MNase samples only when Ca²⁺ was present, confirming site-specific activation of the tethered nuclease. **(D)** Structural localization of MNase cleavage sites on the *Drosophila* 80S ribosome (PDB ID: 6XU8). The cleaved regions on the 18S rRNA (yellow) are highlighted at positions 660 - 672 (green), 1303 - 1312 (magenta), and 1481 - 1492 (blue). The mapping shows that MNase-dependent cleavages cluster within the body and head regions of the 40S subunit.

Mapping the sequencing reads to the *Drosophila* 18S and 28S rRNAs revealed distinct clusters of MNase-dependent cleavage sites that were absent in both the no Ca²⁺ and soluble MNase control samples, confirming that the observed cuts originated from the tethered MNase activity (Figure 5). The cleavage profiles produced by the MNase-FMRP and FMRP-MNase fusions were highly reproducible across biological replicates and, unexpectedly, showed strikingly similar patterns on the 18S rRNA despite the opposite orientations of MNase relative to NT-dFMRP. Both fusions generated prominent cleavage clusters at positions 660-672, 1303-1312, and 1481-1492 within the 18S rRNA (Figure 5 and Supplemental Figure 1). In contrast, cleavage within the 28S rRNA was observed primarily in reactions containing soluble, untethered MNase, indicating non-specific background digestion (Supplemental Figure 2). Notably, the majority of 28S rRNA 3′-end reads mapped to position 1803, which corresponds to a natural break site in *Drosophila* 28S rRNA. This observation further validates the poly(A)-tagging and high-throughput sequencing strategy used to identify MNase-dependent cleavage sites with nucleotide precision.

When the mapped cleavage sites were projected onto the *Drosophila* 80S ribosome structure, the 660-672 cluster localized to the body region, whereas the 1303 - 1312 and 1481 - 1492 clusters mapped to the head region of the 40S subunit (Figure 5D). The 1303 - 1312 cluster aligns with the location of NT-dFMRP in the cryo-EM structure. ^36^ The 660 - 672 cluster is in the expansion segment 6S (ES6S), and the 1481-1492 cluster is in the expansion segment 9S (ES9S), which are highly flexible and possibly readily cleaved by MNase. The absence of substantial cleavage on the 28S rRNA indicates that both termini of NT-dFMRP are oriented toward the solvent-exposed surface of the 40S subunit rather than the intersubunit interface. Notably, the nearly identical cleavage profiles obtained from the N-and C-terminal MNase fusions suggest that both termini of FMRP are highly mobile when bound to the ribosome. The MNase moiety, tethered through flexible linkers at either terminus, appears to explore the surrounding space through rotational and translational diffusion, thereby gaining transient access to multiple exposed regions of the 18S rRNA. This mobility likely reflects an intrinsic conformational flexibility of the N-and C-terminal regions of FMRP, which may undergo dynamic rearrangements relative to the ribosome surface. Consequently, the diffuse rather than sharply localized cleavage pattern supports a model in which NT-dFMRP interacts with the ribosome through a dynamic ensemble of conformations rather than a single static binding mode. These findings highlight the ability of TM-map to capture such conformational heterogeneity in ribonucleoprotein complexes.

## CONCLUSIONS

Traditional methods for studying RBP-ribosome interactions, such as cryo-EM, chemical footprinting, and crosslinking-based approaches, have provided valuable structural and biochemical insights into translation regulation. ^10–16,43^ However, these techniques are often constrained by the need for highly purified and homogeneous complexes, specialized instrumentation, and limited resolution for flexible or transiently bound factors. TM-map complements these approaches by directly coupling RBP-ribosome binding to localized RNA cleavage, thereby converting spatial information into a sequencing-readable signal.

The nucleotide-level precision achieved by TM-map arises from its sequencing-based readout, which captures the exact 3′ termini of MNase-generated fragments. Each cleavage event serves as a positional marker that reflects the local environment of the tethered RBP on the ribosome. Mapping these termini onto ribosomal structures allows the binding site to be inferred within a few angstroms while requiring only standard molecular biology reagents and sequencing workflows. TM-map, therefore, provides an accessible and highly quantitative means to translate spatial proximity into nucleotide-resolution maps of protein-RNA interfaces.

Beyond its resolution, TM-map is inherently scalable and broadly applicable. Distinct RBPs can be profiled in parallel by multiplexed barcoding of sequencing libraries, enabling comparative analyses of multiple factors under identical conditions. Because TM-map relies on genetically encoded MNase fusions rather than large quantities of purified protein, it is well-suited for studying low-abundance or difficult-to-purify RBPs. Importantly, the method is not limited to the ribosome. TM-map can, in principle, be applied to map the interactions of RBPs with other large RNA assemblies such as the spliceosome, signal recognition particle, long noncoding RNAs, or microRNAs. By defining the cleavage signatures generated by individual proteins within these complexes, TM-map could reveal the organization and dynamics of RNA-protein assemblies that are otherwise challenging to characterize structurally.

In addition to identifying binding sites, TM-map can also provide insights into conformational changes within large RNA-protein complexes. Because MNase cleavage is sensitive to the spatial arrangement of the tethered protein and its RNA environment, shifts in cleavage patterns can report structural rearrangements that occur during RNA processing, assembly, or catalysis. Applied over time or under different functional states, the TM-map could thus serve as a powerful reporter of conformational dynamics in multicomponent RNPs.

TM-map is also compatible with *in vivo* applications. ^23–25^ Expression of an MNase-tagged RBP within cells, followed by controlled activation of MNase, would allow mapping of RBP-RNA contacts directly in their native physiological context. Such *in situ* mapping could capture the dynamic association of translation factors, RNA helicases, or regulatory RBPs with ribosomes under stress, nutrient limitation, or developmental transitions. When coupled with transcriptome-wide sequencing, TM-map could reveal how ribonucleoprotein interactions remodel in response to cellular states, providing a bridge between structural and functional genomics.

Looking ahead, TM-map can be integrated with complementary approaches to deepen our understanding of RNA-protein interactions. Coupling TM-map data with cryo-EM reconstructions could guide model fitting or validate flexible and transient factors that elude visualization in density maps. Likewise, integrating TM-map with proximity-labeling techniques such as APEX2 or BioID could link spatial binding information to the identity of associated proteins and RNAs in living cells. ^44^ By bridging high-resolution structural information with transcriptome-scale analyses, TM-map offers a versatile framework for dissecting the dynamic architecture of RNA-protein complexes. Ultimately, this approach promises to illuminate how regulatory RBPs and RNA machineries coordinate gene expression and RNA metabolism across diverse cellular environments.

## ACKNOWLEDGMENTS

This work was supported by the National Institutes of Health (R35GM141864 to S.J.).

## Accession Codes

MS2: P03612

MNase: 1109959A

FMRP: B3LF78

## Contributions

C.Y.Y. and S.J. designed the experiments. C.Y.Y. performed the experiments, and all authors discussed the results. C.Y.Y. and S.J. wrote the paper. S.J. supervised all aspects of the work.

