## Supplemental Figures for "Mapping RNA-Binding Proteins on the Ribosome by Tethered Micrococcal Nuclease"

**Supplemental Figure 1. Non-specific cleavage of *Drosophila* 18S rRNA.**

**(A - D)** Distribution of normalized 3′-termini read counts mapped to selected regions of the *Drosophila* 18S rRNA. Panels show cleavage profiles spanning nucleotides 171 - 180 (A), 471 - 480 (B), 491 - 500 (C), and 1551 - 1560 (D). The x-axis denotes the nucleotide position along the rRNA, and the y-axis represents normalized counts (NC) corresponding to the number of sequencing reads terminating at each position. Read counts were normalized by setting the total for authentic 3′ termini to 100. The error bars indicate the SD from two independent experiments. Cleavage patterns are shown for soluble MNase control (blue bars), MNase-FMRP + $Ca^{2+}$ (red bars), MNase-FMRP without $Ca^{2+}$ (orange bars), FMRP-MNase + $Ca^{2+}$ (green bars), and FMRP-MNase without $Ca^{2+}$ (purple bars). The comparable normalized read counts between the MNase-FMRP and FMRP-MNase reactions and the soluble MNase control indicate that these cleavages are non-specific background events rather than sites of tethered MNase activity.

**Supplemental Figure 2. Non-specific cleavage of Drosophila 28S rRNA.**

**(A - C)** Distribution of normalized 3′-termini read counts mapped to selected regions of the *Drosophila* 28S rRNA. Panels show cleavage profiles spanning nucleotides 1799 - 1808 (A), 2291 - 2300 (B), and 2941 - 2950 (C). The x-axis indicates nucleotide position along the rRNA, and the y-axis represents normalized counts (NC) corresponding to the number of sequencing reads terminating at each position. Read counts were normalized by setting the total for authentic 3′ termini to 100. The error bars indicate the SD from two independent experiments. Cleavage patterns are shown for soluble MNase control (blue bars), MNase-FMRP + $Ca^{2+}$ (red bars), MNase-FMRP without $Ca^{2+}$ (orange bars), FMRP-MNase + $Ca^{2+}$ (green bars), and FMRP-MNase without $Ca^{2+}$ (purple bars). The comparable normalized read counts between the MNase-FMRP and FMRP-MNase reactions and the soluble MNase control indicate that these cleavages are non-specific background events rather than sites of tethered MNase activity. Notably, a strong signal corresponding to the natural break in the 28S rRNA at position 1803 was detected, demonstrating that the poly(A)-tailing, library preparation, and sequencing workflow accurately identifies authentic 3′ termini of rRNA molecules.

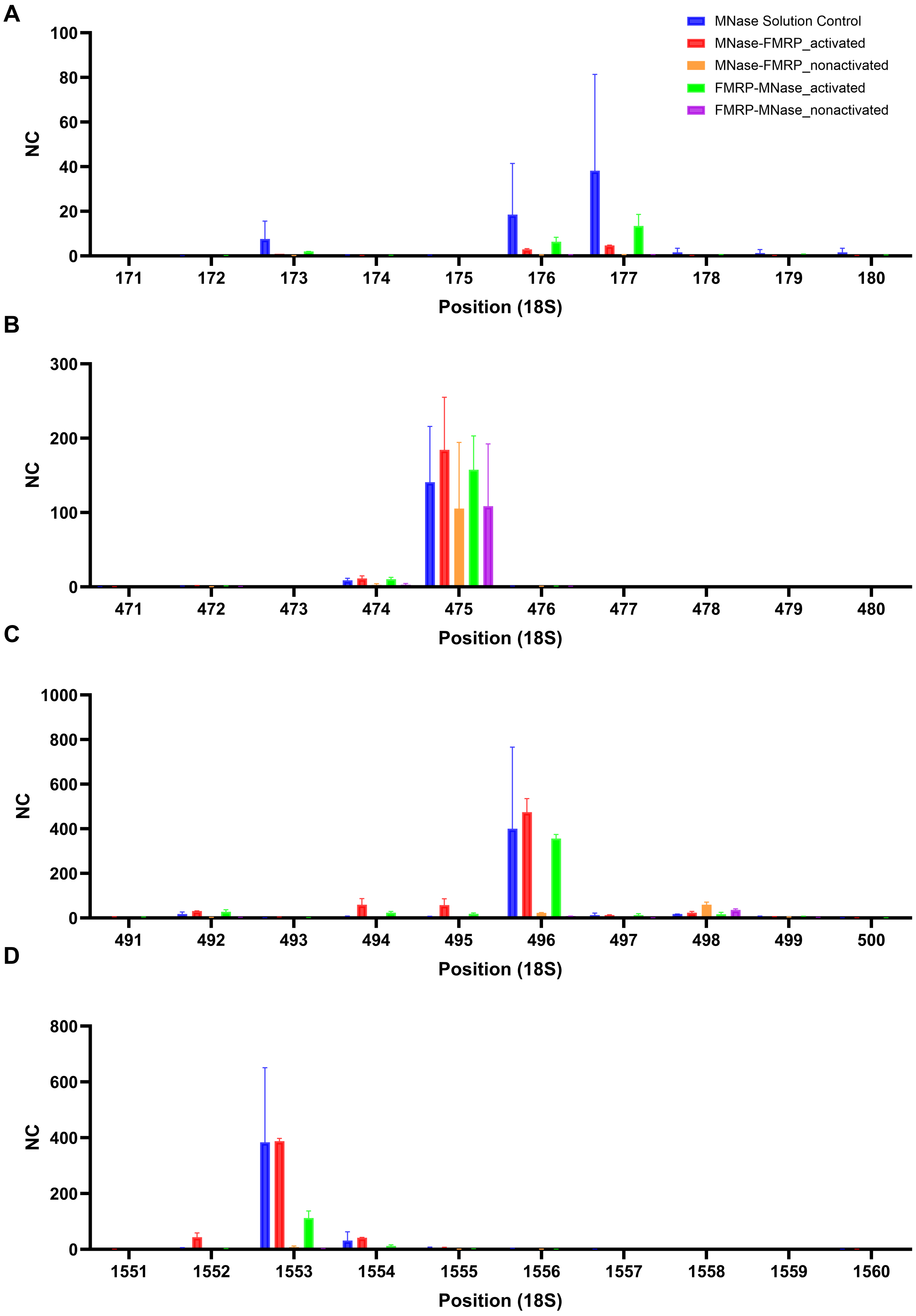


**Supplemental Figure 1**

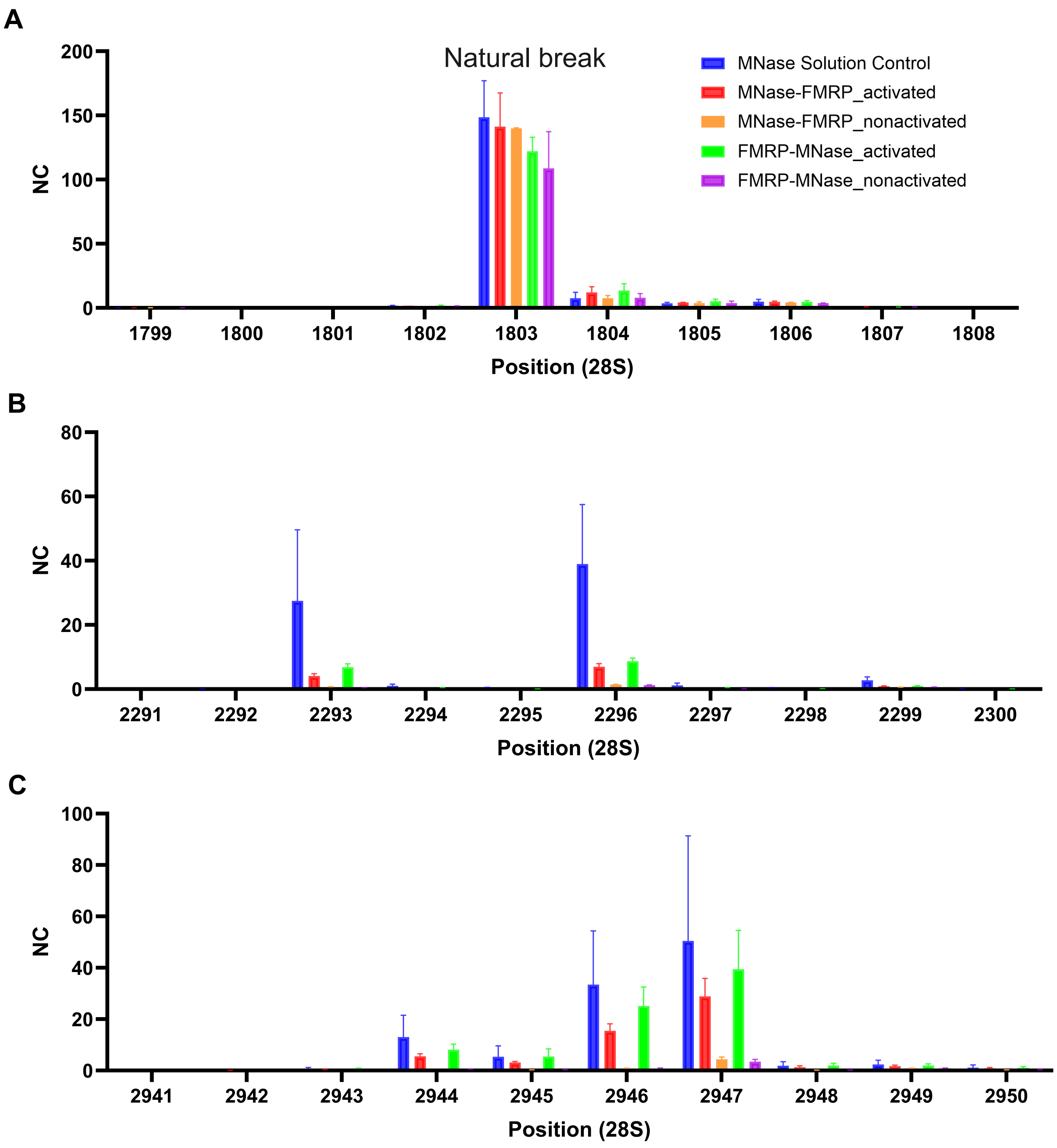


**Supplemental Figure 2**
